# Dynamic regulation of long non-coding RNAs across asexual and gametocyte development in *Plasmodium falciparum*

**DOI:** 10.64898/2026.04.15.718799

**Authors:** Janne Grünebast, Ritwik Singhal, Sophie Olson, Robin Bromley, Sachie Kanatani, Katie Ko, Julie Dunning Hotopp, Photini Sinnis, Manuel Llinás, David Serre

## Abstract

Long non-coding RNAs (lncRNAs) are critical regulators of gene expression in eukaryotes. Short reads from Illumina sequencing, reverse transcriptase artefacts, and incomplete second-strand degradation in strand-specific cDNA libraries hamper genome-wide identification of lncRNAs, especially in gene-dense genomes such as *Plasmodium*. Here, we integrated long-read Oxford Nanopore Technology direct RNA sequencing, ribosome profiling, and single-cell transcriptomics to generate a robust and stage-specific characterization of *P. falciparum* lncRNAs. We generated comprehensive annotations of lncRNAs expressed in both asexual and sexual blood stages and confirmed their non-coding nature using ribosome profiling. Most lncRNAs showed pronounced stage-specific expression and appeared to be particularly abundant in mature gametocytes. Single-cell RNA sequencing revealed differential expression of many lncRNAs in female and male gametocytes, suggesting important roles in gametocytogenesis and transmission. Many lncRNAs are located antisense to protein-coding genes and are co-expressed with their sense mRNA, possibly from putative bidirectional promoters, while others overlap mRNA coding sequences or 3’ untranslated regions and showed negatively correlated expression patterns. Overall, our study shows the prevalence of *P. falciparum* lncRNAs and highlights their possible roles in controlling the regulation of gene expression, particularly during gametocytogenesis.

## Introduction

Malaria still represents a significant global health challenge, with an estimated 282 million new infections in 2024 [1]. *Plasmodium falciparum*, a unicellular protozoan parasite, is the main cause of malaria and is responsible for 95% of malaria cases. It mainly affects individuals from sub-Saharan Africa and results in a high mortality rate, especially in children below the age of five years [1]. The elimination of malaria remains challenging, in part due to increased resistance of the parasites to antimalarial drugs, insecticide resistance of the mosquito vectors, and our incomplete understanding of the complex parasite biology [2]. The parasite is injected into the dermis as sporozoites by a female *Anopheles* mosquito and enters the bloodstream to be carried to the liver, where it infects and multiplies within hepatocytes. Merozoites are released from a liver schizont into the bloodstream, and infect red blood cells (RBCs) [3]. Inside RBCs, the parasite enters the asexual replication cycle. It matures through rings, trophozoites, and schizonts, which again burst and release more merozoites into the bloodstream to infect new RBCs. All malaria symptoms - fever, anemia, headache, chills, and fatigue - arise during this asexual replication, and in severe cases, the infection can lead to cerebral malaria, which often leads to death [1, 2]. A subset of these blood-stage parasites commits to sexual differentiation and develops into female and male gametocytes [4]. These gametocytes are essential for transmission: when taken up by a mosquito blood meal, they fertilize and continue their development in the mosquito, producing the next generation of infectious sporozoites [2, 3].

This complex life cycle is tightly regulated by the expression of ∼5,400 protein-coding genes that are densely packed in 14 nuclear chromosomes, with more than 50% of the genome sequence encoding protein-coding genes and many genes overlapping each other [5]. The parasite has a relatively small number of transcription factors (TFs), including 30 ApiAP2 proteins [6], yet orchestrates highly dynamic and tightly-timed programs to regulate mRNA levels. To achieve this exquisite regulation, additional layers of complexity are likely involved, including epigenetic marks [7–10], post-transcriptional mechanisms, and long non-coding RNAs [11–18].

Long non-coding RNAs (lncRNAs) are emerging as important regulatory elements of gene expression in eukaryotes. LncRNAs are transcribed via RNA Polymerase II, are longer than 200 nucleotides, contain a 5’ cap structure and a poly(A) tail, and can be spliced. A large subset of lncRNAs is transcribed opposite to protein-coding genes and is classified as natural antisense transcripts (NATs) [19]. NATs have been shown to be involved in regulating the transcription of the sense gene, its transcript stability, and/or its translation [19]. A subset of these NATs is transcribed from bidirectional promoters [19]. Antisense RNAs can also recruit histone-modifying proteins or form DNA-RNA triplexes by binding to double-stranded DNA, leading to epigenetic changes that can result in gene silencing or activation [19].

LncRNAs have been described in *P. falciparum* based on analysis of RNA sequencing experiments. One of the best-studied examples is an lncRNA antisense to the gametocyte development 1 (*gdv1*) gene [12]. GDV1 regulates gametocytogenesis by removing Heterochromatin Protein 1 (HP1) from the *ap2-g* locus, the master transcription factor regulating gametocytogenesis [20–23]. Removal of HP1 from *ap2-g* activates the locus, allowing the parasites to undergo sexual commitment and mature into gametocytes [24]. The antisense lncRNA of *gdv1* has been shown to act as a negative regulator of *gdv1,* but its exact mode of action remains uncharacterized and, for example, it is unknown whether it acts through transcriptional interference or by regulating *gdv1* stability or translation [12]. Another well-characterized example is an antisense lncRNA of the Male development protein 1 (*md1*) gene, which is expressed only in female gametocytes, while *md1* is expressed as a male- and female-specific isoform [14]. Another prominent example of lncRNAs in *Plasmodium* are those associated with *var* genes, which encode for *P. falciparum* erythrocyte membrane protein 1 (PfEMP1) [11, 15, 17, 25]. Those intron-derived antisense transcripts are specifically expressed from the active *var* gene locus and play a central role in regulating monoallelic expression, thereby contributing to antigenic variation and parasite immune evasion [13, 26]. A recently identified exonic antisense and downstream intergenic *var*-lncRNAs suggest that a complex network of lncRNAs may regulate the expression of this multi-gene family [17, 27]. In addition to these studies, lncRNAs have also been studied in the context of telomere-associated repetitive elements (TAREs) [28], in relation to their nuclear or cytoplasmic localization, and gametocyte-specific lncRNAs have been identified [16]. Beyond these specific examples, little is known about the overall number of *Plasmodium* lncRNAs, their expression, and their roles, and we still lack a robust and comprehensive annotation of lncRNAs in *P. falciparum*. Previous studies have primarily examined lncRNAs using strand-specific short-read Illumina RNA sequencing [29, 30], which complicates annotations of lncRNAs given the gene-rich nature of the *P. falciparum* genome and the incomplete annotations of the transcripts’ untranslated regions (UTRs). In addition, most RNA sequencing studies rely on cDNA synthesis and can therefore suffer from artefacts introduced during reverse transcription and incomplete strand specificity [16, 29–31]. One recent study manually annotated lncRNAs expressed in mixed asexual *P. falciparum* cultures using Oxford Nanopore Technologies (ONT) direct RNA-sequencing (dRNA-seq) [27, 32]. ONT dRNA-seq has the advantage of sequencing full-length RNA molecules without relying on reverse transcription, enabling robust annotation of lncRNAs.

Here, we use ONT dRNA-seq to characterize and annotate lncRNAs throughout the entire *P. falciparum* asexual development and gametocytogenesis. By comparing ONT data with Illumina short-read RNA-seq, we demonstrate that reverse transcription does lead to spurious antisense transcripts and complicates robust identification of antisense lncRNAs from short-read data. We describe a curated and comprehensive catalog of lncRNAs expressed by blood-stage *P. falciparum* parasites at different stages and validate that they are not translated using ribosome profiling. We further characterize the dynamics of lncRNA expression across the parasite life cycle using single-cell RNA sequencing (scRNA-seq), revealing widespread stage-and sex-specific regulation. Finally, we investigate the relationship between lncRNAs and their neighboring protein-coding genes, uncovering evidence of coordinated transcription from bidirectional promoters and mutually exclusive expression patterns.

## Results

### ONT direct RNA sequencing enables robust strand-specific characterization of RNA molecules

Many lncRNAs are transcribed in the antisense orientation of, and often overlapping with, protein-coding genes, which can complicate their identification after reverse transcription if the strand-specificity is (even partially) lost [33, 34]. For example, commercial stranded (or directional) RNA seq library preparation kits typically use dUTP (deoxyuridine triphosphate) incorporation, instead of dTTP, during second-strand cDNA synthesis and, after adapter ligation, degrade the uracil-containing strand using uracil-DNA glycosylase [35]. However, this enzymatic degradation, while highly efficient, is not perfect, leading to incomplete strand-specificity. In addition, reverse transcription can introduce further artefacts caused by template switching, internal priming, bias towards secondary structures, and incomplete reverse transcription of the mRNAs [34].

To analyze and robustly annotate polyadenylated lncRNAs in *P. falciparum,* we used Oxford Nanopore Technology direct RNA sequencing, the only approach currently able to directly sequence native RNA molecules (and not cDNA). We sequenced RNA isolated from parasites from two different strains cultured *in vitro*. First, we sequenced RNAs isolated from rings, trophozoites, schizonts, and stage V gametocytes from a Malawi field isolate (MWI). Second, we analyzed a tightly synchronized time course of Dd2 parasites and sequenced RNA isolated at 10, 20, 30, and 40 hours post-invasion (hpi), from sexually committed rings, and from stage I-V gametocytes. In total, we generated 11,768,245 reads across 14 samples, with 575,655-1,411,388 reads per sample.

We first compared the ONT data with Illumina short-read RNA-seq data generated from the same RNA sample to assess strand-specificity in both datasets, focusing on reads spanning splice sites of protein-coding genes, for which antisense reads are most likely spurious (see Materials and Methods). In Illumina RNA-seq, 0-59% of these spliced reads (mean 1.46%) mapped in antisense orientation (Fig. 1a), confirming the incomplete strand-specificity. By contrast, no spliced ONT reads were mapped on the antisense strand of a specific exon-exon junction (Fig. 1b). While the proportion of spurious antisense reads introduced by incomplete digestion of dUTP is, on average, relatively small, this artefact qualitatively hinders the accurate annotation of antisense transcripts.

**Fig. 1:**
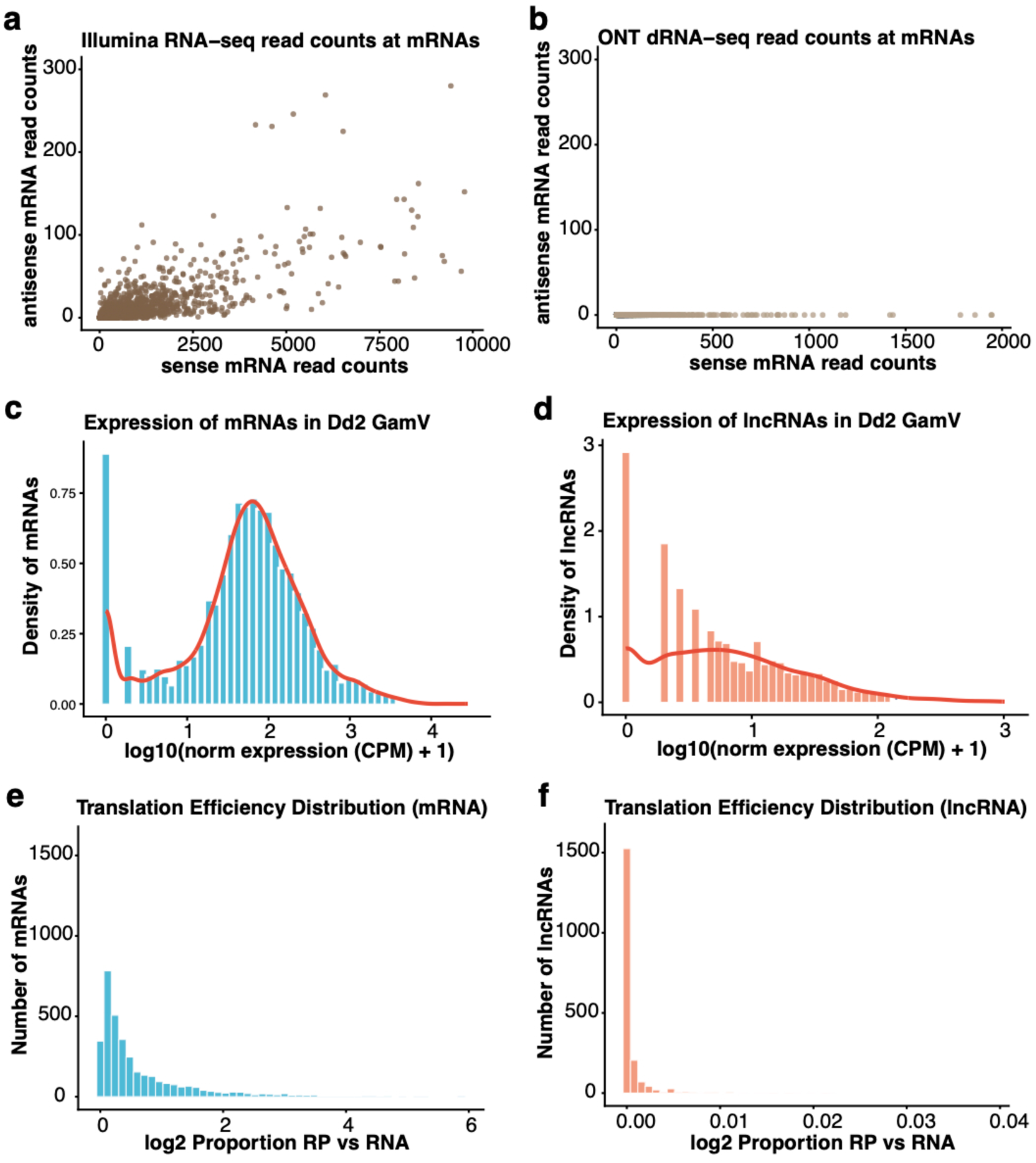
General features of lncRNA and mRNA. Strand specificity of Illumina RNA-seq **(a)** and ONT direct RNA sequencing (dRNA-seq) **(b)** assessed by counting reads spanning splice junctions of protein-coding genes on sense (x-axis) and antisense strands (y-axis). Antisense splice-junction reads represent reverse-transcription artefacts. Distribution of the expression levels of mRNAs **(c)** and lncRNAs **(d)** in Dd2 stage V gametocytes (based on the ONT dRNA-seq data). Translation efficiency of mRNAs **(e)** and lncRNAs **(f)** in NF54 asexual parasites, calculated as the ratio of normalized ribosome profiling reads to normalized RNA-seq reads per transcript.

### Characterization of P. falciparum lncRNAs using ONT dRNA-seq and Ribosome Profiling

We used the 14 ONT sequencing datasets generated from the Dd2 and MWI parasite strains to annotate long non-coding RNAs in the asexual and sexual blood stages of the parasite. 60-91% of the reads in each sample mapped to annotated protein-coding genes, while 4-33% mapped to rRNAs (see Materials and Methods for details). The remaining of the reads were further analyzed and curated to remove reads potentially derived from fragmented mRNAs and incompletely annotated protein-coding genes. Overall, out of 7,186 - 46,780 reads overlapping lncRNAs, we annotated 2,186 putative lncRNAs that did not encode for any amino acid sequences and were longer than 200 nucleotides.

On average, these lncRNAs were 1,518 bp nucleotide long (vs. 3,023 for annotated protein-coding genes), with extensive variations (ranging from 200 bp to 7,858 bp), and around 30% of lncRNAs contained multiple exons. lncRNAs were also typically expressed at a lower level than mRNAs. For example, in Dd2 stage V gametocytes, the mean normalized expression for lncRNAs was 20 cpm (count per million reads), while mRNAs were expressed at an average of 180 cpm (Fig. 1 c & d). Note, however, that lncRNA expression levels varied extensively and that some lncRNAs were highly expressed. For example, a lncRNA transcribed near RAD54, a protein important for DNA repair [36], showed an expression of 235 cpm in stage Dd2 stage V gametocytes, much higher than the expression of RAD54 with 145 cpm, while another lncRNA, transcribed near P25, an important ookinete surface protein [37], was also highly expressed (199 cpm in Dd2 stage V gametocytes).

To confirm that the lncRNAs identified were indeed non-coding, we performed ribosome profiling on a mixed asexual NF54 parasite culture. We identified RNAs bound to active ribosomes and compared translation signals at mRNAs and lncRNAs. While the proportion of mRNAs bound to ribosomes varied depending on the gene (Fig. 1e), ribosome occupancy at lncRNAs was consistently much lower, if at all detectable (Fig. 1f). In addition, most of the lncRNAs associated with ribosomes displayed unequal occupancy throughout their sequence (as we would expect for translation of a potential open reading frame), possibly indicating their involvement in the regulation of translation (for example, some lncRNAs have been shown in other eukaryotes to interact with ribosomes or act as ribosome sponges [38]).

### Stage- and sex-specific expression of lncRNAs

To preliminarily assess how mRNAs and lncRNAs differ among developmental stages, we first performed principal component analysis (PCA) using, separately, the expression of all annotated protein-coding gene mRNAs and lncRNAs. For both mRNAs and lncRNAs, the two strains cluster together at each developmental stage, while the expression profiles of each stage recapitulated their relative position in the life cycle (e.g., PC1 separating samples along gametocytogenesis).

To gain a better understanding of the variations in lncRNA expression during development, we then focused on the ONT data generated from the tightly synchronized cultures of Dd2 parasites. Out of 797 lncRNAs highly expressed in Dd2 parasites (i.e., ≥20 cpm in at least one stage), 491 (61%) were most abundantly expressed in stage I - V gametocytes (Fig. 2a), with most lncRNAs displaying the highest expression in stage V gametocytes (n=240). Of the lncRNAs most expressed during asexual replication, most lncRNAs were expressed in schizonts (n=196). We also identified 57 lncRNAs with the highest expression in sexually committed rings. These findings highlight extensive stage-dependent regulation of lncRNAs.

**Fig. 2:**
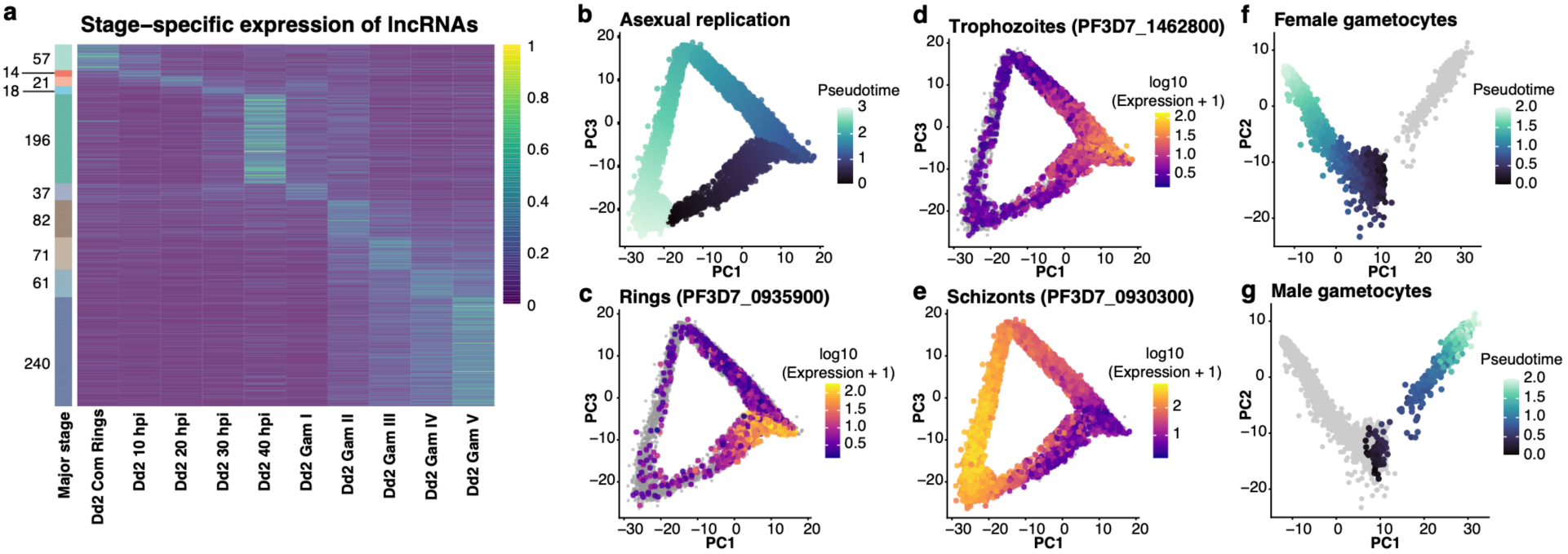
Stage-specific lncRNA expression and single-cell RNA-seq analysis. **(a)** Heatmap of strand-specific lncRNA expression across Dd2 blood stages from ONT dRNA-seq data, colored according to their expression level (in blue to green scale). The stacked bar on the left shows the number of lncRNAs most expressed in a given stage. **(b-e)** PCA of individual NF54 parasites characterized by scRNA-seq: asexual parasites **(b)** and female **(f)** and male gametocytes **(g)** colored according to their pseudotime **(b)** or to the expression of marker genes (rings: PF3D7_0935900 **(c)**, trophozoites: PF3D7_1462800 **(d)**, and schizonts: PF3D7_0930300 **(e)**.

Even though the cultures analyzed by ONT dRNA-seq were highly synchronized, the bulk approach might mask subtle variations in development or, in the case of the gametocyte cultures, differences between male and female development. To further investigate stage- and sex-specific expression of lncRNAs, we generated scRNA-seq datasets from both i) an asynchronous asexual culture and ii) mature gametocyte cultures of NF54 parasites. After stringent QC, we retained 14,604 asexual parasites, capturing the entire intraerythrocytic replication cycle from rings to trophozoites to schizonts (Fig. 2b-e), and 7,133 gametocytes, recapitulating the female and male gametocyte development (Fig. 2f,d).

Since scRNA-seq experiments rely on reverse transcriptase, these data may be affected by incomplete strand-specificity and complicate analyses of antisense transcripts (see above). For all further analyses, we therefore (when possible) specifically analyzed the subsets of scRNA-seq reads spanning exon-exon junctions of a given mRNA or lncRNA to prevent artefacts. Because lncRNAs are generally lowly expressed and scRNA-seq only captures relatively highly expressed transcripts, only a subset of lncRNAs (and mRNAs) could be reliably analyzed. Among these, 95% of lncRNAs were expressed exclusively in gametocytes (n = 103). Of these gametocyte-expressed lncRNAs, 48.5% were detected only in female gametocytes, 38.8% only in male gametocytes, and 12.6% in both sexes. In contrast, protein-coding gene mRNAs were often expressed throughout multiple consecutive blood stages.

The widespread sex-specific expression of lncRNAs suggests that these transcripts may play important roles in gametocyte differentiation and function. To further explore this observation, we analyzed several representative lncRNAs in detail, highlighting previously characterized examples as well as newly identified sex-specific transcripts. One of the lncRNAs showing sex-specific expression is a previously characterized lncRNA expressed antisense to the *male development 1* (*MD1*) gene [14]. Consistent with previous report, the scRNA-seq data shows that i) *md1* has four exons and is transcribed into multiple isoforms that are expressed in male and female gametocytes, with the exception of the isoform containing exons 1 and 2 that is exclusively detected in male gametocytes (Fig. 3a, left and middle panels), while the antisense lncRNA is exclusively expressed in female gametocytes (Fig. 3a, right panel). In addition to the previously described lncRNA, we also observed a second, asexual-specific, lncRNA antisense of *md1* (Fig. 3a, schematics, grey).

**Fig. 3:**
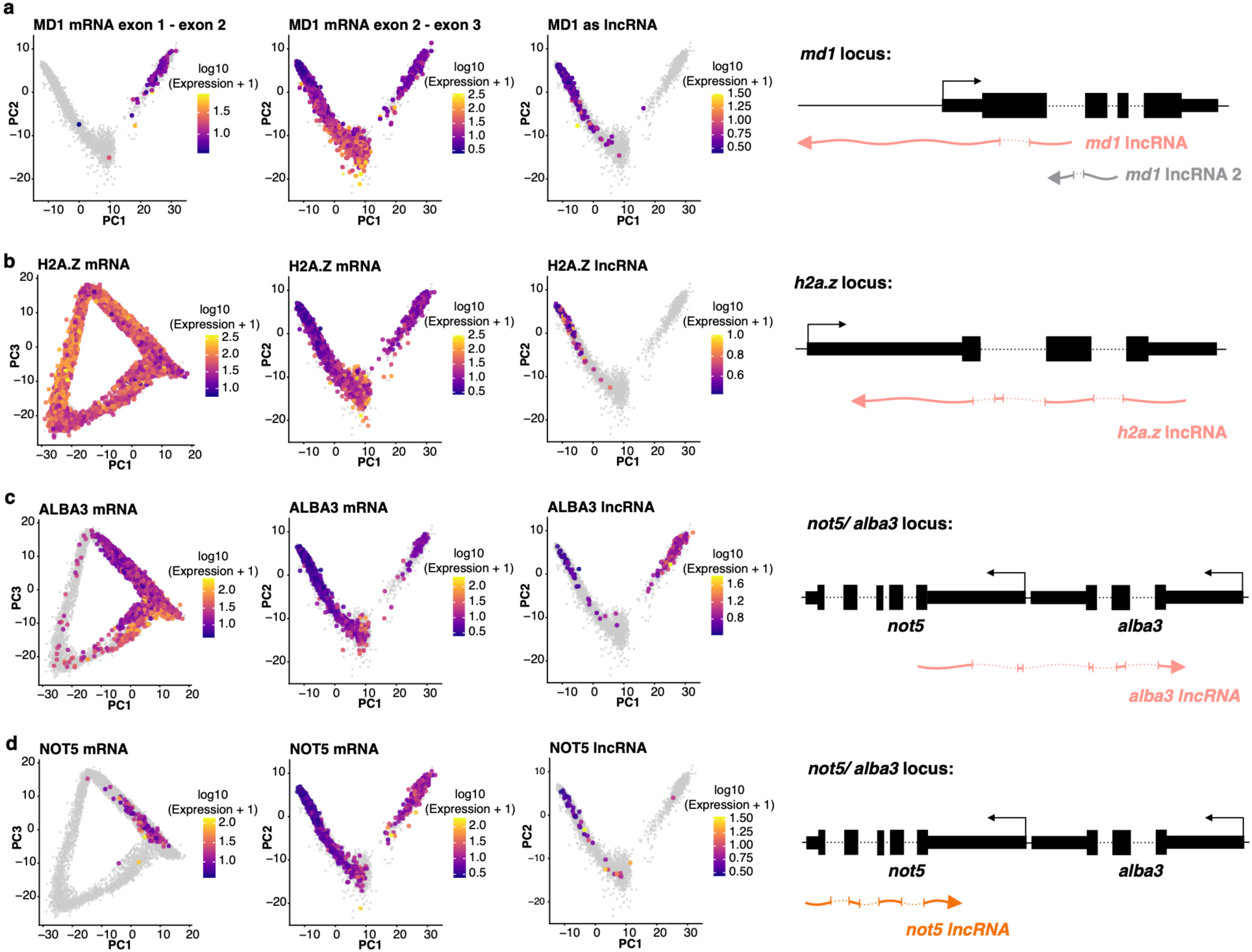
Sex-specific lncRNA expression in gametocytes. **(a)** Expression of *MD1* mRNA at exon 1-2 (left) and exon 2-3 splice junctions (middle) in NF54 gametocytes, with antisense lncRNA spliced junction reads (right). A schematic of the *MD1* locus illustrates the gene structure (black) and the antisense lncRNA orientation (pink), including a second asexual-specific lncRNA (grey). **(b-d)** Expression of mRNAs (*H2A.Z*, *ALBA3*, *NOT5*) along asexual and gametocyte pseudotimes.

Beyond this known example, we identified many new sex-specific lncRNAs. For example, an antisense lncRNA of *H2A.Z*, an alternative histone variant enriched in euchromatin regions during the asexual replication [39], showed female gametocyte-specific expression, whereas the *H2A.Z* mRNA was expressed throughout the parasite life cycle with the highest expression during the asexual replication and early gametocytogenesis (Fig. 3b). One lncRNA overlapping both the 3’UTR/CDS of *ALBA3*, which is potentially involved in the DNA damage response [40], and the 5’UTR of *NOT5*, a component of the CCR4-NOT involved in mRNA turnover [41], was exclusively expressed in mature male and female gametocytes with higher expression in males (Fig. 3c) while the *ALBA3* and NOT5 protein-coding transcripts were expressed more broadly, with the highest expression in asexual ring and trophozoite stages (Fig. 3c & 3d). Note that a second lncRNA antisense to the 3’UTR and CDS of *NOT5* was also detected but expressed only in female gametocytes and at very low levels (Fig. 3d). Together, these findings illustrate the complexity of gene expression in *Plasmodium*, with many distinct lncRNAs expressed in stage- and sex-specific manner and distinctly from the sense mRNAs.

### Genomic location and expression of lncRNAs with regard to protein-coding genes

We then examined the locations of the highly expressed lncRNAs (≥20 cpm) with regard to neighboring mRNA expression of protein-coding genes (Fig. 4a). Due to the conservative criteria chosen for our identification pipeline, putative lncRNA sequences overlapping and in the same direction as an annotated protein-coding transcript were often disregarded as likely fragmented mRNAs. As a consequence, most of the newly identified lncRNAs are natural antisense transcripts (NATs, n=1685) or intergenic (i.e., not overlapping with any annotated protein-coding genes, n= 655). NATs were further divided into divergent and convergent lncRNAs, based on the overlap with the sense protein-coding gene (following the nomenclature described in [19]). 726 lncRNAs located antisense to a gene and with a transcription start site within the 5’ UTR of the sense transcript (‘overlapping’) or within 1,000 bp of an annotated 5’ UTR (‘nonoverlapping’) were classified as divergent NATs (Fig. 4a & 4b). LncRNAs being transcribed antisense to a gene with the transcription initiation site either downstream of or within the gene were broadly categorized as convergent NATs (Fig. 4a & 4b, n = 959). Overall, these analyses indicated that most newly identified lncRNAs were positioned antisense or intergenic relative to protein-coding genes, with a substantial fraction classified as divergent or convergent NATs.

**Fig. 4:**
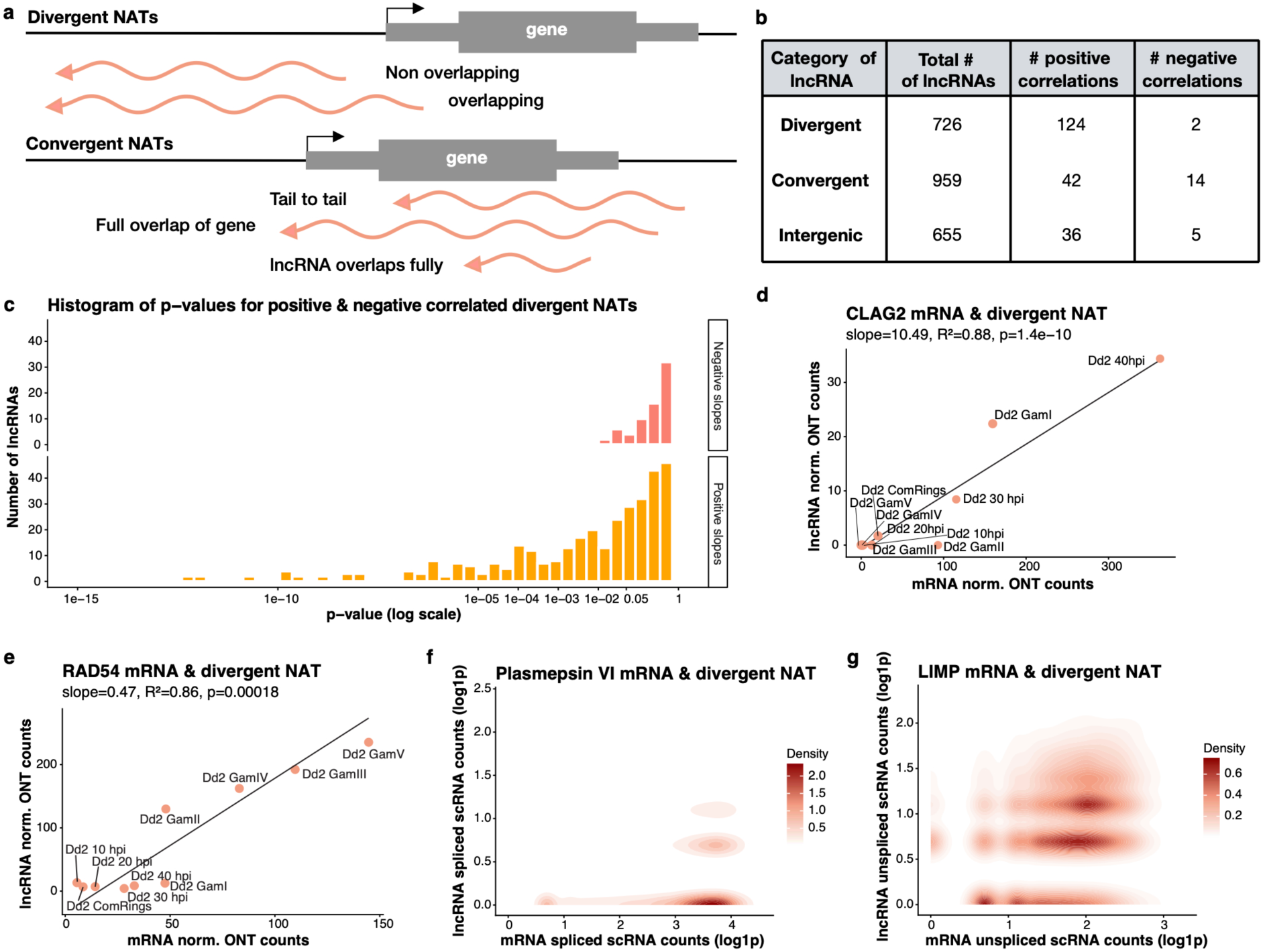
Correlations between lncRNA and mRNA expression and bidirectional promoters. **(a)** Schematic showing the different categories of natural antisense transcripts (NATs). Divergent NATs arise from overlapping 5′ UTRs or from non-overlapping regions that start within 1 kb of a protein-coding gene. Convergent NATs involve tail-to-tail overlap of 3′ UTRs, lncRNAs spanning an entire gene, or lncRNAs fully embedded within a gene. **(b)** Number of lncRNAs in each category and a summary of the correlation analysis between the expression of lncRNAs and neighboring mRNAs using robust linear regression. **(c)** Histogram of p-values for all positive and negative correlations between NATs and neighboring mRNAs, including non-significant pairs, separated by correlation direction (positive vs. negative slope). **(d,e)** Example of the correlations between mRNA expression (x-axis) and lncRNA expression (y-axis) determined by ONT dRNA-seq from Dd2 parasites at different stages at the *CLAG2* **(d)** and *RAD45* **(e)** loci. **(f,g)** Examples of correlations between mRNA expression (x-axis) and lncRNA expression (y-axis) in individual parasites as determined by scRNA-seq at the *Plasmepsin VI* **(f)** and *LIMP* **(g)** loci (only spliced are counted to avoid RT artefacts at the *Plasmepsin VI* locus, while since the transcripts do not overlap, unspliced reads were considered for *LIMP*).

We then tested whether the lncRNA expression was statistically correlated with the mRNA expression of neighboring genes across stages using ONT dRNA-seq data from the tightly synchronized Dd2 cultures. In total, we identified 202 significantly positively correlated mRNA/lncRNA expression pairs and 21 negatively correlated pairs (p ≤ 0.01) (Fig. 4b-e). Interestingly, among the significant pairs, the expression of divergent NATs was almost always (98%) positively correlated with the expression of the sense transcript (Fig. 4b & 4c), while 25% of the convergent NATs were negatively correlated with the expression of the neighboring sense gene.

### Bidirectional promoters and divergent lncRNAs

The positive correlation between the expression of many divergent NATs and their neighboring genes (see, e.g., Fig. 4d & 4e) could indicate the presence of a bidirectional promoter that simultaneously regulates the expression of both loci. Alternatively, since the dRNA-seq data is a bulk approach, the positive correlations observed do not necessarily represent co-expression within a single cell but instead reflect that some cells expressed the lncRNAs and some other cells expressed the neighboring mRNAs, leading to a spurious statistical association. To evaluate this possibility, we examined, at single-cell resolution, whether the mRNA and divergent NAT were transcribed in the same parasite. The low lncRNA abundance (Fig. 1d) complicated genome-wide assessment of lncRNA expression from scRNA-seq data, but a few divergent NATs were sufficiently expressed to support these analyses. For example, the expression of a lncRNA at the *plasmepsin IV* locus, was only detected in cells also expressing the protein-coding gene, an aspartic protease important for degrading hemoglobin within the food vacuole [42] (Fig. 4f). Similarly, the expression of the lncRNA associated with *LIMP*, a gliding-associates protein translationally repressed in gametocytes [43], was correlated among cells with the expression of the protein-coding gene (Fig. 4g). These results clearly support the hypothesis that divergent NATs are co-expressed with their neighboring gene, likely from a putative bidirectional promoter.

One possible explanation for this co-expression from a bidirectional promoter is that transcription is “leaky” and that the lncRNA is generated as a byproduct of mRNA transcription, but without having any functional relevance. Alternatively, the shared promoter could indicate a genuine coordinated regulation, with both the mRNA and lncRNA being functional and possibly participating in related biological pathways. Two lines of evidence argue against the leaky promoter hypothesis. First, only a small proportion (17%) of all NAT/mRNA pairs were significantly correlated, suggesting that this is not an overall feature of *Plasmodium* promoters but a genuine pattern specific to some lncRNAs. Second, while the mRNAs are typically expressed at higher levels than the lncRNAs (Fig. 1c,d and Fig. 4d for an example), in some cases the lncRNA is expressed at a higher level than its neighboring mRNA. For example, the expression of the lncRNA overlapping the 5’UTR of *RAD54*, a putative DNA repair protein, is correlated with RAD54 mRNA expression but the lncRNA is actually more abundant than the mRNA (Fig. 4e). This observation contradicts the hypothesis that the promoter is leaky and transcription of the lncRNA is just a by-product, and instead support the co-expression of the mRNA and lncRNA, possibly because they regulate different part of the same process.

### Convergent lncRNAs may interfere with the expression of neighboring mRNAs

In contrast to the high number of divergent lncRNAs whose expression was positively correlated with the expression of the neighboring mRNAs, several convergent lncRNAs appeared to be negatively correlated with the expression of the neighboring mRNA. One example is the translation initiation factor eIF-1A, which has an antisense lncRNA overlapping the 3’ UTR and part of the CDS (Fig. 5a). Translation initiation factor eIF-1A is involved in initiating translation by scanning mRNAs for the start codon, and together with eIF1, it maintains an open ribosome conformation [44]. Based on the ONT dRNA-seq data, the *eIF-1A* mRNA was expressed in all stages, with variable expression levels, and seemed to display an opposite expression than its overlapping lncRNA: *eIF-1A* mRNA seemed lowly expressed in stage II-V gametocytes, where we observed the highest expression of its antisense lncRNA (Fig. 5b). We further investigated the expression patterns through scRNA-seq data and observed that *eIF-1A* mRNA is expressed throughout the life cycle, with lower expression in schizonts (Fig 5c,d). In contrast, the *eIF-1A* lncRNA is mainly expressed in female gametocytes (Fig. 5e,f). LncRNAs can act at the post-transcriptional level by influencing mRNA stability and regulating translation [19], and we therefore wanted to know whether the lncRNA and mRNA were expressed in the same cell. scRNA-seq data revealed that, overall, the mRNA and lncRNA were typically not expressed in the same parasites (Fig. 5g-i). This expression pattern suggests a possible regulatory relationship between the antisense lncRNA and the *eIF-1A* mRNA, possibly indicating that the lncRNA is negatively regulating mRNA abundance during gametocyte development (the few cells displaying expression of both the mRNA and lncRNA could further indicate that this regulation occurs post-transcriptionally (e.g., by affecting the mRNA stability rather than inhibiting transcription) but further studies will be required to characterized the molecular mechanisms underlying this regulation).

**Fig. 5:**
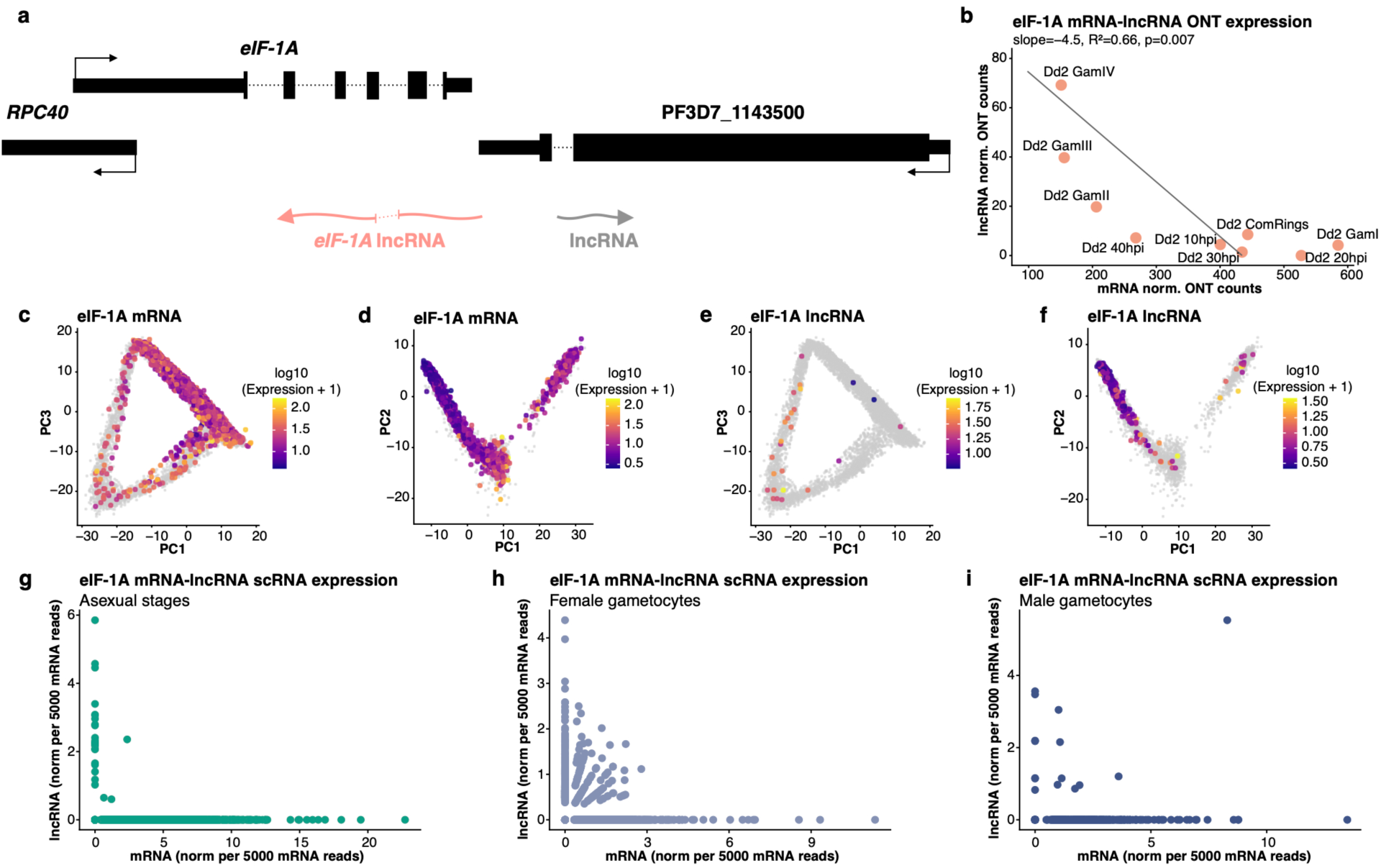
Opposite expression of lncRNA and mRNA at the *eIF-1A* locus. **(a)** Schematic of the lncRNA at the *eIF-1A* locus. **(b)** Correlations between eIF-1A mRNA expression (x-axis) and lncRNA expression (y-axis) determined by ONT dRNA-seq across stages. **(c, d)** Expression of *eIF-1A* mRNA in asexual **(c)** and gametocyte **(d)** using spliced scRNA-seq reads. **(e,f)** Expression of the antisense *eIF-1A* lncRNA in asexual **(e)** and gametocyte **(f)** using spliced scRNA-seq reads. **(e-g)** Correlations between mRNA expression (x-axis) and lncRNA expression (y-axis) in individual parasites as determined by scRNA-seq in **(e)** asexual stages, **(f)** female gametocytes, and **(g)** male gametocytes; each dot represents a single cell.

## Discussion

Long non-coding RNAs are emerging as critical regulators of eukaryotic gene expression, yet their roles in *P. falciparum* have only started to be unraveled [11–14, 17, 28]. The genome of *P. falciparum* is gene-dense, with 50% of its sequence encoding proteins, overlapping genes, incompletely annotated UTRs and antisense transcription, making it challenging to accurately annotate lncRNAs from short-read Illumina sequencing data. Here, we leveraged Oxford Nanopore Technologies’ direct RNA sequencing to generate a high-confidence catalog of lncRNAs expressed across asexual and sexual stages of *P. falciparum*. ONT dRNA-seq allows direct sequencing of native RNA molecules, which circumvents biases introduced by reverse transcriptase and short-read libraries. Specifically, mispriming during reverse transcriptase, template switching, and incomplete strand specificity can artificially lead to natural antisense transcripts. By comparing long-read ONT sequencing and Illumina short-read RNA-seq, we confirmed that ONT provides a complete strand separation of transcripts. This enabled generating a robust and comprehensive annotation of lncRNAs in the *P. falciparum* genome, particularly for those antisense of protein-coding genes. Ribosome profiling data confirmed that the vast majority of the newly annotated lncRNAs were not bound to ribosomes and are indeed non-coding.

Bulk ONT dRNA-seq across tightly synchronized Dd2 parasites revealed that lncRNAs are highly stage-specific, with distinct lncRNAs expressed during asexual replication, sexual commitment, and gametocytogenesis. Within the asexual cycle, most lncRNAs are predominantly expressed in schizonts and sexually committed rings, suggesting potential roles in cell cycle progression and the initiation of sexual differentiation. Late-stage gametocytes show the highest abundance of lncRNAs, suggesting a specific role for non-coding transcription prior to transmission. Because our bulk ONT data do not allow distinguishing male from female gametocytes, we also integrated single-cell RNA-seq data. These data enabled the analysis of transcriptional heterogeneity at the level of individual cells and the examination of sex-specific lncRNA expression patterns. Many lncRNAs showed specific expression in either male or female gametocytes (and a few were expressed in both sexes). We speculate that this high number of lncRNAs present in female gametocytes could be involved in the regulation of translational repression since, in humans, lncRNAs have been shown to interact with RNA-binding proteins that control stress granule formation [45].

We confirmed the sex-specific expression of a previously characterized lncRNA, antisense of MD1 [14]. In addition, many lncRNAs show sex-specific expression. One particularly interesting lncRNA was expressed exclusively in female gametocytes antisense of H2A.Z, a histone variant usually incorporated at promoter regions in euchromatic regions during asexual replication [39]. *H2A.Z* is highly expressed throughout the asexual life cycle and in both male and female gametocytes. Previous studies have shown that in female gametocytes, H2A.Z is enriched in H3K9me3-associated heterochromatic regions, in addition to being enriched in promoter regions [10]. Given the established role of H2A.Z in heterochromatin organization in female gametocytes, this lncRNA could represent an additional layer of control over chromatin state and gene expression. Similarly, we identified an antisense lncRNA that overlapped both the 3’ UTR and part of the coding sequence of *ALBA3*, and extended into the promoter region and 5’ UTR of *NOT5*. NOT5 is a component of the CCR4-NOT complex, a key regulator of mRNA turnover in eukaryotes. In *Plasmodium yoelii,* components of this complex, including the CCR4-1 deadenylase, have been implicated in regulating transcript abundance and coordinating male gametocyte maturation [41]. ALBA3 is an apurinic/apyrimidinic (AP) endonuclease with potential roles in the DNA damage response and has been reported to inhibit transcription by binding to DNA [40, 46]. While *ALBA3* is expressed during asexual stages, particularly in rings and trophozoites, and at lower levels in gametocytes, expression of the associated lncRNA was restricted to gametocytes and showed higher expression in males. A second lncRNA, antisense of *NOT5*, was lowly expressed in female gametocytes. *NOT5* is very lowly expressed in trophozoites, but is expressed in both female and male gametocytes, with a higher expression in males. This divergence in expression patterns suggested that these lncRNAs were not simply a byproduct of *ALBA3* and *NOT5* transcription (see also below) but may instead modulate their activity through transcriptional or post-transcriptional mechanisms. Expression of sex-specific lncRNAs may contribute to the divergence of male and female developmental programs, which are essential for successful transmission. In addition, this layered regulation is particularly relevant given the extensive translational repression observed in gametocytes, raising the possibility that lncRNAs help inhibiting translation and/or enhancing the stability of these transcripts that are only translated after transmission to the mosquito.

Analysis of the expression of lncRNAs and their neighboring mRNA revealed the complexity and multifaceted impacts of lncRNAs on gene expression. Expression of many divergent NATs showed a positive correlation with the expression of the adjacent mRNA, possibly indicating their coordinated regulation from a bidirectional promoter. Single-cell analyses further demonstrated that these divergent NATs and their corresponding mRNAs were frequently co-expressed within the same cell, supporting their coordinated transcription (rather than a statistical artefact arising from averaging independent processes occurring in different cells). Bidirectional promoters have previously been described between two protein-coding genes and between a protein-coding gene and NATs [31, 47, 48]. In addition, the use of plasmid systems with bidirectional promoters is well established in *P. falciparum* [49]. One explanation for this pattern is that these lncRNAs are a byproduct of leaky promoters. However, only a fraction of all divergent NATs showed such a strong correlation, suggesting that most lncRNAs are not functionally irrelevant transcripts generated spuriously by a “leaky” transcription. This byproduct hypothesis is further contradicted by instances, such as the *RAD54* locus, where the lncRNA is more expressed than the mRNA.

In contrast, we observe that the expression of some convergent lncRNAs, overlapping the 3’UTR and/or the CDS of a sense mRNA, was negatively correlated with the expression of their sense mRNA. One striking example is an antisense lncRNA of the translation initiation factor *eIF-1A*, which showed stage-specific expression in gametocytes, whereas *eIF-1A* mRNA was largely absent in the same cells (with a few cells in female gametocytes where both lncRNA and mRNA were expressed). Notably, *eIF-1A* mRNA levels were reduced at stages when the lncRNA was most abundant, suggesting that the lncRNA could negatively regulate mRNA expression, potentially through a post-transcriptional regulatory mechanism. This putative regulation of eIF-1A is particularly interesting in the context of the translational repression of mature gametocytes [50–52].

By integrating long-read sequencing, ribosome profiling, and single-cell analysis, we provide a robust resource and a solid foundation to systematically characterize the role of lncRNAs in the regulation of *Plasmodium* gene expression. These genome-wide analyses also provide some new hypotheses of the possible role of these lncRNAs and suggest that they might participate in the regulation of gene expression by affecting transcription, through transcriptional interference or binding to DNA and modifying the chromatin conformation, mRNA stability, or decay. In this regard, it is fascinating that we observed most lncRNAs expressed in late gametocytes when translation is inhibited and many mRNAs are stored in Ribonucleoprotein (RNP) stress granules. One could speculate that some of these lncRNAs are key regulators of this process by stabilizing mRNAs [53], regulating translation (e.g., by binding to the translation start site or translation initiation factors [54–56]), or by participating in the RNP complexes [45, 57]. Further studies, leveraging the information generated here, will need to rigorously examine these mechanisms and decipher this additional level of complexity of *Plasmodium* gene expression regulation.

## Methods

### Parasite culture

Parasites were cultured at 3% hematocrit and synchronized twice using 5% sorbitol. For Dd2, asexual parasites were collected at 10, 20, 30, and 40 hpi (hours post-invasion) and stored in TRIzol LS (Thermo Fisher) for RNA extractions. Gametocytes of Dd2 and NF54 parasites were induced as previously described in [58]. In brief, 20-24 hpi parasites, initially grown in incomplete RPMI media supplemented with 25 mM HEPES (pH 7.4), L-glutamine, 0.2% sodium bicarbonate, 0.1 mM hypoxanthine, 50 μg/mL gentamycin, and 0.25% AlbuMAX® (complete medium), were diluted to 1.5-2% parasitemia in 25-50 mL of minimal fatty acid (mFA) medium. The mFA medium consisted of incomplete RPMI with 0.39% fatty-acid free BSA, 30 μM oleic acid, and 30 μM palmitic acid. At 20-22 hours post-induction (day 1), the medium was replaced with complete RPMI, and from day 2 onward, 20 U/mL heparin was added daily. Sexually committed Dd2 rings were collected at 30 hours post mFA induction, while gametocytes from Dd2 were harvested on days 2, 4, 6, 8 and 12 and stored in TRIzol LS. NF54 gametocytes for ribosome profiling and scRNA-seq experiments were harvested at day 16. In addition, asynchronous NF54 parasites were used for ribosome profiling and scRNA-seq. *P. falciparum* IGS-MWI, a clinical isolate from Malawi, was cultured at the University of Maryland, Baltimore. Incomplete RPMI supplemented with 25 mM HEPES, L-glutamine, 0.24% sodium bicarbonate, 0.367 mM hypoxanthine, 25 µg/mL gentamycin, 0.25% AlbuMAX®, and 5% heat-inactivated human serum was used, and parasites were cultured at 4% hematocrit. Asexual stages were synchronized with sorbitol, harvested at rings, trophozoites, and schizonts (confirmed by microscopy), lysed with saponin, and resuspended in 500 µL TRIzol LS (Thermo Fisher). Gametocytes were induced using the “crash” method [59]. Cultures were synchronized at 3-5% ring-stage parasitemia (day 0) with 5% sorbitol. On day 1, spent media was replaced with pre-warmed incomplete media containing 10% human serum. On day 2, cultures were diluted to 0.1% rings, and media was changed daily without adding red blood cells until stage V gametocytes were harvested on day 16. Gametocytes were enriched using a multilayer Percoll gradient (80%, 65%, 50%, 35%) and collected from the 35/50% interface, lysed with saponin, and resuspended in 500 µL TRIzol LS (Thermo Fisher). RNA was extracted as previously described using chloroform and ethanol precipitation [58].

### Oxford Nanopore direct RNA sequencing

ONT dRNA-seq was performed using the *P. falciparum* Dd2 strain and the IGS-MWI strain. 500 ng of RNA was used per sample, and the libraries were prepared using the Direct RNA Sequencing library preparation kit (SQK-RNA002). Each sample was sequenced using a FLO-MIN106D Spot-ON Flow Cell (R9 Version) for 72 hours.

### Bulk RNA-sequencing using Illumina short-reads

150 ng of RNA from *P. falciparum* IGS-MWI parasites was used for selecting polyadenylated sequences using the NEBNext® Poly(A) mRNA Magnetic Isolation Module, and RNA libraries were generated using the NEBNext® Ultra™ II Directional RNA Library Preparation Kit following the protocol. The quality of the libraries was checked using the Agilent TapeStation system, and sequencing was performed on a NovaSeq 6000 with 150 bp paired-end reads.

### Ribosome Profiling

Ribosome profiling was performed using the ALL-IN-ONE RiboLace Gel Free kit from Immagina Biotechnology [60]. An asynchronous asexual culture of *P. falciparum* NF54 parasites was used. Red blood cells were lysed with 0.1% saponin and washed 3 times with 1x PBS. The parasites were lysed in the supplemented lysis buffer and incubated on ice for 30 min. The parasites were stroked using a pre-chilled douncer homogenizer for 300 strokes, with a 30-second pause every 30 strokes. The lysed parasites were centrifuged at 20,000 x g for 5 min to separate the nuclei, and the supernatant was transferred to a new tube. This parasite lysate was stored at -80 °C until further processing. From here, the protocol was followed. In short, beads were functionalized with the Rsp probe, and successful probe binding was confirmed using the Nanodrop. The cell lysate was digested with Nuclease (Nux) for 45 min at 25 °C, and the digestion was stopped by adding SUPERaseIn (Thermo Fischer) for 10 min on ice. This digested cell lysate was then incubated with the functionalized beads for 70 min at 4 °C. RNA was extracted and used for library preparation. First, the 5’ end was phosphorylated, followed by adapter ligation and circularization. After this reverse transcription, two PCR amplifications were performed. The quality of the library was assessed using the Agilent Tapestation system with a D1000 DNA Screentape. For the parasite lysate, RNA was extracted using TRIzol LS (Thermo Fisher) as previously described [58], and RNA-seq libraries were generated using the NEBNext® Poly(A) mRNA Magnetic Isolation Module and the NEBNext® Ultra™ II Directional RNA Library Preparation Kit (NEB). The libraries were sequenced on a NovaSeq 6000 using 150 bp paired-end sequencing.

### Single-cell RNA sequencing (10X Genomics)

Single-cell RNA sequencing was performed using a *P. falciparum* NF54 asynchronous asexual culture and a mature-stage V gametocyte culture. Infected red blood cells were purified using MACS columns, and 20,000 cells per sample were used for library preparation using the Chromium GEM-X Single Cell 3’ Kit v4 (10X Genomics), following the manufacturer’s instructions. The quality of the libraries was checked using an Agilent TapeStation system, and sequencing was performed on a NovaSeq 6000 with 150 bp paired-end sequencing.

### Annotation of lncRNA from ONT dRNA-sequencing data

ONT data from *P. falciparum* Dd2 and MWI parasites were used to annotate lncRNAs. ONT reads were basecalled using a GPU Guppy version 6.4.2 (--config rna_r9.4.1_70bps_hac.cfg --min_qscore 7 --records_per_fastq 10000000 --gpu_runners_per_device 8 --num_callers 1). The FASTQ file was then aligned against the *P. falciparum* 3D7 reference genome from PlasmoDB (version 63) using minimap2 [61] with the following options: -ax map-ont -t 2. SAM files were sorted and indexed using samtools version 1.20 [62], and the quality of the reads was visualized using IGV version 2.16.0. Using custom scripts for the annotation of lncRNAs, reads overlapping ≥50% with annotated protein-coding genes or pseudogenes were saved in a new BAM file, and stringtie version 2.2.1 [63] was used to extend the annotation of protein-coding genes. Next, the reads that overlapped <50% in the first step (potential non-coding) were further assigned as putative coding (≥50%) or putative non-coding (<50%) based on the new stringtie annotation. From the putative non-coding BAM file, reads overlapping snoRNAs, tRNAs, and rRNAs were filtered out. Stringtie was run on this filtered non-coding file, and the annotation with the highest coverage was kept. Transdecoder Version 5.7.1 was used to check for potential open reading frames in this non-coding annotation, and transcripts with complete open reading frames encoding proteins ≥100 amino acids were discarded. In addition, partial open reading frame predictions were filtered out when the putative non-coding RNA overlaps with a known protein-coding gene. This led to a putative non-coding BAM file for each stage and a predicted annotation from Stringtie, which was manually corrected with WebApollo and IGV to annotate lncRNAs supported by at least five reads across all stages. This annotation was again filtered for tRNAs, rRNAs, and snoRNAs, and Transdecoder was used to identify complete open reading frames to filter out. All non-coding RNAs < 200 base pairs were filtered out, and the lncRNAs were renamed and saved as a new annotation in a GTF file.

### Bioinformatics analysis of sense- antisense counts from RNA-seq and ONT

Illumina RNA-seq data from MWI stage V gametocytes were aligned against the reference genome of *P. falciparum* 3D7 (v63) using HiSat version 2.2.1 [64], with both paired-end sequencing files (R1 and R2) and a maximum intron length of 5000. Samtools version 1.20 was used to process SAM files and sort them into BAM files. ONT reads were processed as described above. Splice-aware RNA-seq analysis was performed by identifying reads spanning introns based on CIGAR strings containing “N” from the BAM files. Exon annotations from protein-coding genes from the reference GFF of the *P. falciparum* PlasmoDB annotation were used to reconstruct gene-specific splice junctions, and only reads with introns matching annotated junctions were retained. PCR duplicates were removed (Illumina data only) based on genomic coordinates, and reads were assigned to genes in a strand-specific manner as sense or antisense. Gene-level counts of correctly spliced reads were generated for the sense and antisense strands, and the percentage of antisense read counts was calculated.

### Bioinformatics analysis of ONT dRNA-seq data

Custom scripts were used to count expression levels based on the PlasmoDB annotation (v63) GFF file and the final lncRNA GTF annotation from the ONT dRNA-seq data. For this, reads overlapping ≥50% and on the same strand as an annotated protein-coding gene were counted towards the gene, and reads overlapping ≥50% and on the same strand as an annotated lncRNA were counted towards the lncRNA. ONT reads were normalized by counts per million (cpm) based on the number of the total number of lncRNA vs mRNA reads.

To assign lncRNAs into different categories, lncRNAs were extracted from the GTF annotation file, and their genomic coordinates, strand orientation, lengths, and exon counts were recorded. Protein-coding gene annotations, including exon structures and untranslated regions (5′ and 3′ UTRs), were obtained from the PlasmoDB reference GFF file and organized by chromosome. For each lncRNA, distances to all mRNAs on the same chromosome were calculated, and the two closest genes were identified. Overlap lengths and percentages between lncRNAs and candidate genes were computed, and strand orientation was used to classify each pair as sense or antisense. Finally, lncRNA-mRNA relationships were categorized into genomic context classes based on relative position, transcription start sites, and overlap with coding or UTR regions. In particular, for antisense-oriented lncRNA-mRNA pairs, the lncRNA was categorized as i) convergent if the lncRNA transcription start site (TTS) was located in the 3’ UTR/CDS or less than 1kb after the end of a gene, overlapping the gene or if the lncRNA fully overlapping with a gene ii) divergent if the lncRNA TSS was located in the 5’ UTR or less that 1kb upstream of the gene (Fig. 4a) or iii) intergenic if the lncRNA was outside of any gene.

To determine the stage at which each lncRNA was most highly expressed, we used normalized read counts across the ten Dd2 samples. For each lncRNA, expression values were filtered to only analyze lncRNAs with a maximum stage-specific expression greater than 20 cpm. The total expression across all stages was calculated, and the fractional expression per stage was derived by dividing individual stage counts by the total expression. The stage contributing to the highest fraction of expression was defined as the major stage for each lncRNA.

### Bioinformatics analysis of ribosome profiling data

Ribosome profiling reads were processed from raw FASTQ files using custom scripts. For this, we only used Read 1 (R 1) sequences containing the expected 3 ′ adapter (TCTCCTTGCATAATCACCAACC), which was subsequently trimmed. Reads shorter than 30 nucleotides after adapter removal were discarded. Unique molecular identifiers (UMIs) were extracted from the first and last four nucleotides of each read, appended to the read header, and removed from the sequence, followed by trimming of a leading thymidine. Processed reads were converted to FASTA format for downstream alignment using HiSat2 Version 2.2.1 with --max-intronlen 5000. SAM files were sorted and indexed using samtools version 1.20 and saved as a BAM file. The quality of the data was checked using IGV. Aligned reads from BAM files were filtered to retain primary and secondary alignments, and PCR duplicates were removed based on genomic position, strand, and UMI using custom scripts. Reads were assigned to genomic features (mRNA, rRNA, tRNA, and lncRNA) using midpoint coordinates of the read and strand-specific overlap with annotated regions from PlasmoDB GFF annotation and the lncRNA GTF annotation. Read counts were calculated per gene (i.e., including mRNAs, lncRNAs, and rRNAs), and summary statistics, including mapping categories and read length distributions, were generated. For RNA-seq data, aligned paired-end reads were similarly processed using custom scripts. For RNA-seq data, paired-end reads (R1 and R2) were aligned using HISAT2 (v2.2.1) with a maximum intron length of 5000. Aligned reads were processed using custom scripts, where read midpoints were calculated based on strand orientation and PCR duplicates were removed using genomic coordinates. Reads were assigned in a strand-specific manner to protein-coding genes and lncRNAs based on overlap with annotated regions, and gene-level counts along with summary statistics were generated. Ribosome profiling and RNA-seq counts were normalized to cpm. Genes with low RNA expression (cpm < 20) were excluded. Translation efficiency (TE) was calculated as the ratio of ribosome-protected fragment abundance to RNA abundance and log2-transformed (log2(TE + 1)). The overall distribution of translation efficiency was compared between the two gene categories.

### Bioinformatics analysis of scRNA-seq data

Single-cell RNA sequencing libraries were processed using custom scripts. Paired-end Illumina reads were used, with Read1 containing the 10X Genomics cell barcode and unique molecular identifier (UMI) and Read2 containing the cDNA sequence. Read1 sequences shorter than 28 nucleotides were discarded. Read2 sequences were trimmed to remove the 5′ template-switching oligo (TSO), 3′ Illumina adapters, and poly-A tails, with reads shorter than 41 nucleotides after trimming removed. Processed reads were converted to FASTA format and aligned to the *Plasmodium falciparum* 3D7 genome using HISAT2 (v2.2.1) with a maximum intron length of 5,000. BAM files were filtered to retain primary alignments, and PCR duplicates were removed based on genomic position, strand, cell barcode, and UMI. Unique reads were assigned to mRNA or rRNA genes based on midpoint read coordinates that overlapped annotated regions in the PlasmoDB GFF annotation. For each cell barcode, per gene counts were recorded. In addition, summary statistics, including mapped reads, PCR duplicates, and unassigned reads, were saved. Cells were then filtered based on total counts of unique reads (i.e., after removing PCR duplicates), and cells with less than 500 unique reads were discarded. We performed single-cell PCA and pseudotime analysis separately for the asexual and the gametocyte datasets using top variable genes (top 500 by variance). Raw counts were first filtered to remove likely doublets (>15,000 unique reads per cell) and then normalized using CPM. PCA was applied on the filtered, normalized counts to visualize major sources of variation. Screen plots were generated to assess the variance explained by each principal component. Marker genes specific to female (PF3D7_1031000) and male (PF3D7_1469900) lineages were normalized per cell (CP5K) and overlaid on the PCA, allowing identification of marker expression per cell. Outliers along PC2 were filtered manually to refine the PCA representations for each sex. Pseudotimes were calculated by measuring the Euclidean distance between cells based on their principal component 1, 2, and 3.

Single-cell RNA-seq data from asexual and gametocyte stages were processed to quantify lncRNA and mRNA expression per cell. Asexual cells were assigned to Ring, Trophozoite, or Schizont stages using marker gene expression, while gametocytes were assigned as Female or Male. Spliced counts for lncRNAs and mRNAs were normalized and averaged per stage. Genes expressed in ≥2% of cells in any stage were retained, and each gene was assigned a “Major Stage” based on the highest fraction of expressing cells. Expression in each stage was determined if lncRNA or mRNA was expressed in at least 2% of cells per stage.

## Acknowledgments

This study was supported by awards from the National Institutes of Health (R21 AI194421 to D.S. and M.L, R01 AI172827 to D.S., U19 AI110820 to D.S. and J.D.H., R01 AI132359 to P.S., and R03AI180804 to S.K.), from the Deutsche Forschungsgemeinschaft (DFG, German Research Foundation - 549182383 to J.G.), from Bloomberg Philanthropies (to P.S. and S.K.), and from the National Science Foundation to J.D.H. (EF 2025384). The funders had no role in study design, data collection and analysis, decision to publish, or preparation of the manuscript.

